# AlphaFind: Discover structure similarity across the entire known proteome

**DOI:** 10.1101/2024.02.15.580465

**Authors:** David Procházka, Terézia Slanináková, Jaroslav Olha, Adrián Rošinec, Katarína Grešová, Miriama Jánošová, Jakub Čillík, Jana Porubská, Radka Svobodová, Vlastislav Dohnal, Matej Antol

## Abstract

AlphaFind is a web-based search engine that provides fast structure-based retrieval in the entire set of AlphaFold DB structures. Unlike other protein processing tools, AlphaFind is focused entirely on tertiary structure, automatically extracting the main 3D features of each protein chain and using a machine learning model to find the most similar structures. This indexing approach and the 3D feature extraction method used by AlphaFind have both demonstrated remarkable scalability to large datasets as well as to large protein structures. The web application itself has been designed with a focus on clarity and ease of use. The searcher accepts any valid Uniprot ID, PDB ID or gene symbol as input, and returns a set of similar protein chains from AlphaFold DB, including various similarity metrics between the query and each of the retrieved results. In addition to the main search functionality, the application provides 3D visualizations of protein structure superpositions in order to allow researchers to instantly analyze the structural similarity of the retrieved results. The AlphaFind web application is available online for free and without any registration at https://alphafind.fi.muni.cz.

**GRAPHICAL ABSTRACT:** 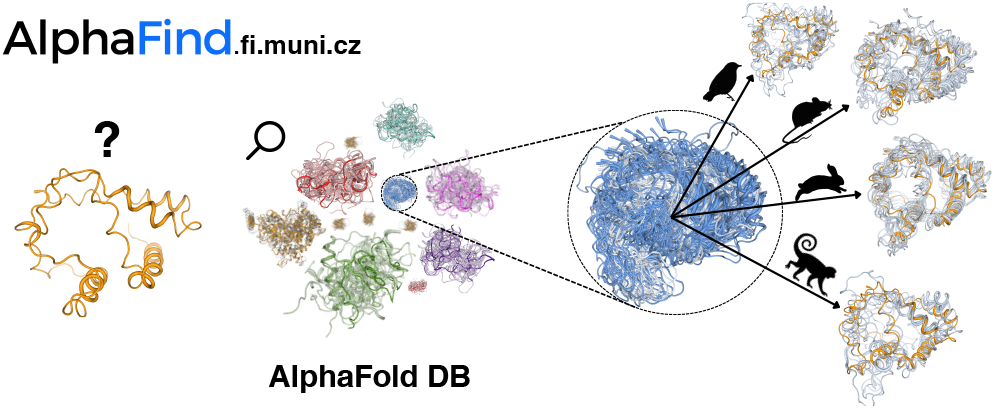

## INTRODUCTION

Protein structural data are highly beneficial scientific resources that serve as the basis for ever-growing and valuable research. Thanks to seven decades of intensive research, we currently have more than 200 thousand experimental protein 3D structures deposited in the Protein Data Bank (PDB) (1). Furthermore, the AlphaFold algorithm (2), trained on these experimental data, produces highly reliable protein 3D structure predictions. As a result, the AlphaFold database has been published (3), containing more than 200 million 3D protein structures. In parallel, other databases of predicted protein 3D structures have been published, such as ESM Metagenomic Atlas (4), collecting 600 million protein 3D structures. Various research fields in bioinformatics benefit from such databases, with new cutting-edge technology on the horizon (5).

To take full advantage of this enormous amount of data, it is essential to organize them efficiently. Specifically, it is necessary to perform a structure-based search, because structural similarities frequently imply functional correspondence, even without high sequence similarity.

Unfortunately, convetional protein structure tools (e.g., PDBeFold) are not able to handle such huge datasets. To address this issue, novel searching tools have been developed, e.g., FoldSeek (6), 3D-surfer (7)or Dali server (8).However their functionality has some substantial limitations: they cannot search through the whole AlphaFold DB, and they rely on predefined fold patterns.

For this reason, we developed AlphaFind, a protein search application that is focused entirely on tertiary structure, automatically extracting the main 3D features of each protein chain and using a machine learning model to find the most similar structures. Therefore, it is not limited by the occurrence of specific fold patterns. AlphaFind takes advantage of the Learned Metric Index approach (9) as well as a 3D feature extraction method designed for protein structures (10),both of which have demonstrated remarkable scalability to large datasets and to large protein structures.

The AlphaFind web application has been designed with a focus on clarity and ease of use, with links to AlphaFold DB provided for additional information wherever appropriate. The search accepts any valid Uniprot ID or PDB ID as input and returns a set of similar protein chains from AlphaFold DB, including various similarity metrics between the query and each of the retrieved results. Beyond the primary search functionality, the application provides 3D visualizations of protein structure superpositions, and allows researchers to instantly analyze the structural similarity of the retrieved results.

## DESCRIPTION OF THE WEB SERVER

The application is available as an online service free of charge. It is composed of two integral components – a front end and a back end. The front end is accessible via an internet browser, communicating with the back end that evaluates queries and returns results back to the front end. The back end starts with an offline phase, where it pre-processes the data and creates a search index for rapid search request evaluation in the online phase.

### Protein structure representation

We utilize the compact data embedding method described in (10) in conjunction with data clustering and machine learning. This approach captures the semantic relationships between protein structures and quickly identifies the most relevant groups of data for a given query. The embedding of a protein is essentially an extremely compressed representation of its 3D structure, which only accounts for the broad structural features of the whole structure while neglecting primary or secondary structure information.

### Data preparation/indexation

In the offline phase, we first extract semantic information from raw *cif* files into vector embeddings, thereby compressing the original AlphaFold DB size from *∼* 23 TiB into *∼* 20 GiB and obtaining representation that is more suitable for data processing algorithms. Then, we build an index on the embeddings using a learned indexing approach. Learned indexes form a new research stream defined by Kraska et al. for one-dimensional numeric data (11).This research direction was later expanded to the domain of complex data where data items are compared using a distance function. The distance expresses complex similarity beyond mere comparison of two integers, as shown in (9, 12). The latter two methods in conjunction with (10) establish the basis of the indexing solution presented in here.

### Workflow

AlphaFind was developed to provide an intuitive and useful interface for discovering structurally similar proteins within AlphaFold DB. The workflow, as depicted in Figure 1, was optimized and tested for maximum computational efficiency to provide fast and accurate results. The following steps take place from the moment that the user poses a query:

**Figure 1.**
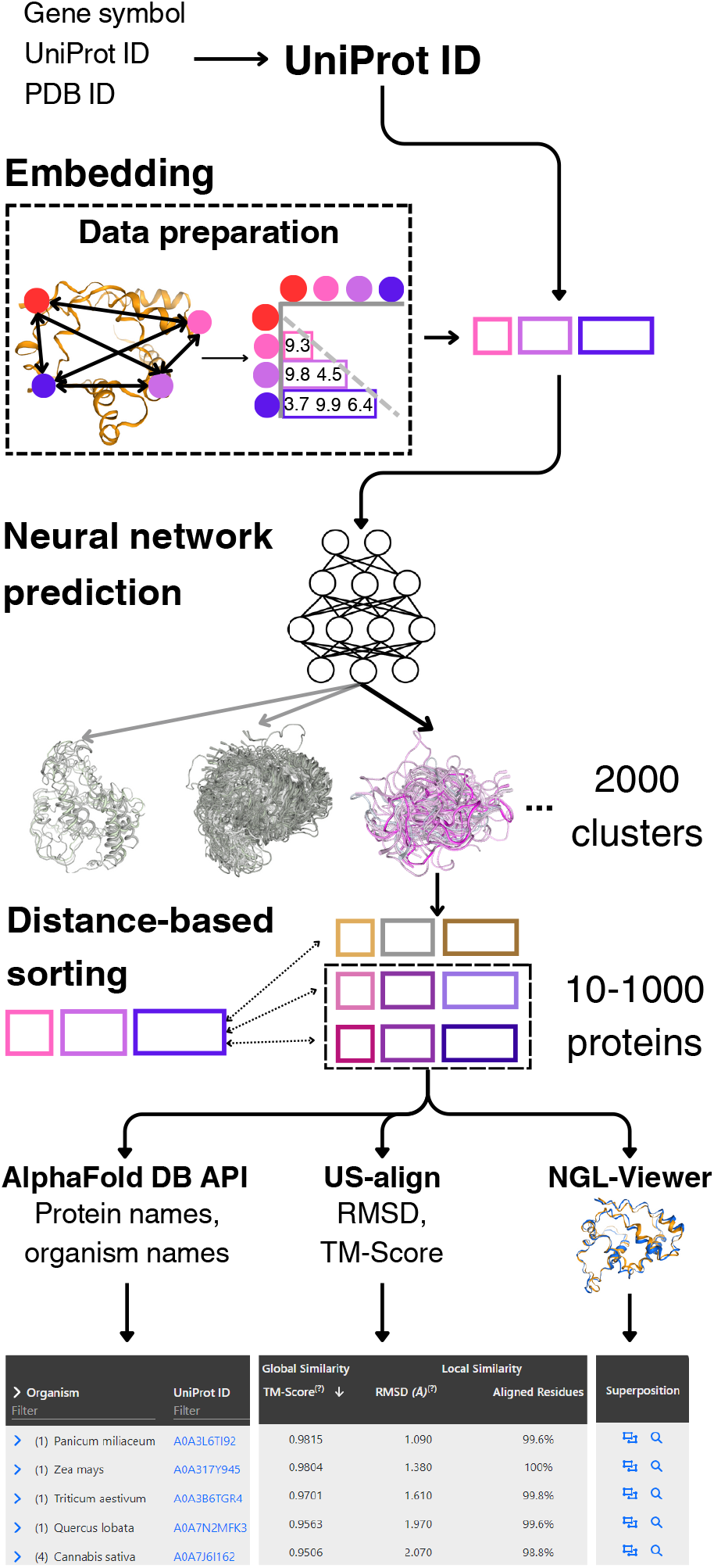
Visual representation of AlphaFind’s workflow.

1. *Translating the input into a UniProt ID*: AlphaFind supports three forms of input: UniProt ID, PBD ID, and Gene symbol. Since UniProt ID is internally used to identify a protein, other forms of input must be translated into UniProt ID using publicly available APIs. For PDB ID to UniProt ID conversion, we use: https://www.ebi.ac.uk/pdbe/api/mappings/uniprot/ and Gene symbol to UniProt ID conversion AlphaFind relies on: https://rest.uniprot.org/idmapping.
2. *Searching for a set of candidate proteins*: using the UniProt ID, the server finds the associated protein structure embedding created during data preparation. This embedding is served to the index (a fully connected neural network), which returns the top ten most similar clusters to the embedding, narrowing down the set of candidates from 214 to *∼*10 million. Then, Euclidean distance is evaluated between the input and each candidate object embedding (13). We sort such candidates by this proximity and take just 1000 closest proteins, which reduces the search space even further.
3. *Evaluating global and local similarity:* Based on the desired result set size (*n* = 50, initially), the nearest *n* proteins out of the candidate set are selected to evaluate local (*RMSD*) and global (*TM-Score*) similarity, together with *aligned residues* and *sequence identity*, using the US-align tool (14). This is the most time-intensive step of the search process, taking up to several tens of seconds.
4. *Downloading metadata for query and results*: After collecting the search results, AlphaFind uses the AlphaFold DB’s API to collect useful information, such as the name of the protein structure, the host organism name, and groups the results by organism name for easier orientation.
5. *Visualizing protein structures’ overlap:* To represent the protein structure similarity visually, AlphaFind utilizes NGL Viewer (15) to display an overlay of every pair of the input protein and a result protein. Additionally, for a more detailed view, each result points to the Mol* Viewer (16).
6. *Showing more results:* The user has the option to expand the search results by additional candidate proteins identified in Step 2: they can choose to display 50, 100, 200, or 300 more. If this option is selected, Steps 3-5 repeat, and the result table is updated.
7. *Downloading the results:* Once searching is finished, the user can download the results for future use or archival in the CSV format using “Export all to CSV” button.

To reduce response times, once Steps 2 and 3 are executed for a given input protein, the web server stores the computed results, causing a significant reduction in subsequent waiting times when searching for the same protein.

### Implementation

The front end was implemented in TypeScript using the React framework (https://reactjs.org/) and the Bootstrap library (https://getbootstrap.com/). The back end is written in Python 3.10 with Flask (https://flask.palletsprojects.com/).

We tested AlphaFind on a diverse set of proteins varying in size, complexity, and quality. AlphaFind provided biologically relevant results even for small, large and lower quality structures. When AlphaFind did not offer structures with high TM-scores, the results remained biologically relevant. A performance of AlphaFind showed as scalable.

### Limitations

AlphaFind is constructed on AlphaFold DB in version 3, predating the v4 update. The application returns up to 1000 similar protein structures to a single query. The results returned by AlphaFind are approximate, and as such, they may not necessarily contain all the most structurally similar proteins present in AlphaFold DB.

Approximate searching trades off resource demands (such as hardware, time and money) for precision. It is customary in complex data management, as the “objective answer”, e.g., which protein structure is more similar to another, does not have an indisputable answer in the first place. Moreover, the design of the application allows for loading additional data, which expands and also refines the prior search answer. This functionality enables the user to find protein structures, which could have been missed in the previously shown search results. AlphaFind processes the entire protein structure – its helices, sheets and its unstructured regions – and handles them with equal weight. Therefore, high occurrence of unstructured regions in the input structure can bias the search. This phenomenon is more prevalent in coiled-coil structures but can be also observed in some small structures.

## RESULTS AND DISCUSSION

### Searching characteristics

Due to the data embedding method used, AlphaFind is mainly concerned with global structural similarity, always comparing the entire structure and treating all parts with equal weight, regardless of primary or secondary structure. Unlike established searching methods, which tend to place more focus on occurrence of certain folds or high local similarity in particular regions, AlphaFind is more likely to find structural similarity across organisms where these regions are less conserved.

AlphaFind indexes the entire AlphaFold DB, which contains 214 million protein structures. The conversion to embeddings allows the application to search the database and return the first 50 results in an average of 7 seconds with negligible back-end load. The user can expand the obtained results in increments of 50, 100, 200, and 300, which takes, on average, additional 7, 9, 11, and 15 seconds, respectively. During periods of high load, the users’ requests may be queued and processed with a slight delay, increasing with the number of concurrent users that interact with the application. To demonstrate AlphaFind in practice, we provide three use cases of search and consequent analysis of the returned results.

Hemoglobin is a protein that facilitates the transport of oxygen and other gases in red blood cells. Almost all vertebrates contain hemoglobin. It consists of four protein subunits (globins), and is one of the first proteins whose 3D structure has been experimentally determined. There are many types of hemoglobin, with hemoglobin Alpha 1 (encoded by the HBA1 gene) occurring in humans and being the main form of hemoglobin in adults. Here, AlphaFind shows us (Figure 2) that highly similar hemoglobin structures can also be found in other species.

**Figure 2.**
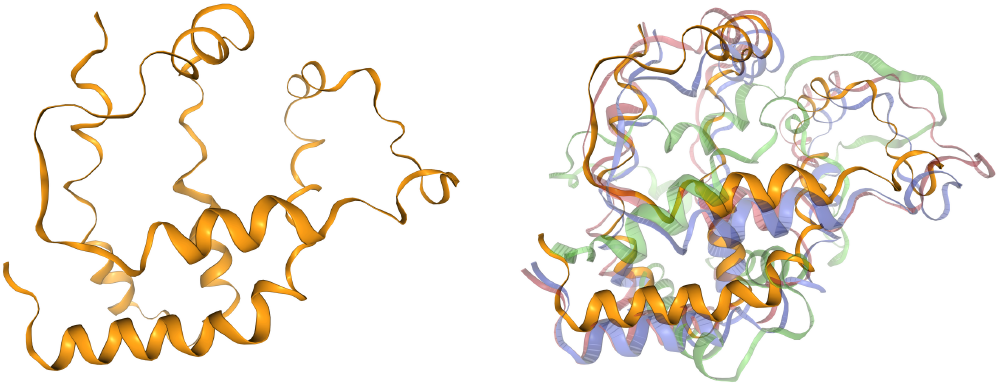
Hemoglobin alpha 1 from *Homo sapiens* (UniProt ID: I1VZV6) in yellow. Examples of AlphaFind search results: Alpha-globin from *Mus musculus* (UniProt ID: Q61287) in blue, Hemoglobin subunit beta-like from *Esox lucius* (UniProt ID: A0A3P8Z522) in green, Hemoglobin alpha A subunit from *Anas georgica* (UniProt ID: B9VY01) in red.

Cytochrome P450 are enzymes which are important for the metabolism of many endogenous compounds and xenobiotics. P450 enzymes have been identified across all biological kingdoms: animals, plants, fungi, bacteria and archaea, as well as in viruses. Cytochrome P450 proteins contain one chain which is composed of more than 20 sheets and helices. Their sequence similarity is very low. In this use case, we can observe similarities among cytochrome P450 structures from various species (Figure 3). The search starts with a cytochrome from corn (*Zea Mays*), and within the first 50 hits, we find similar structures originating from various animals (fish, eagle, mouse, cat, horse, etc.).

**Figure 3.**
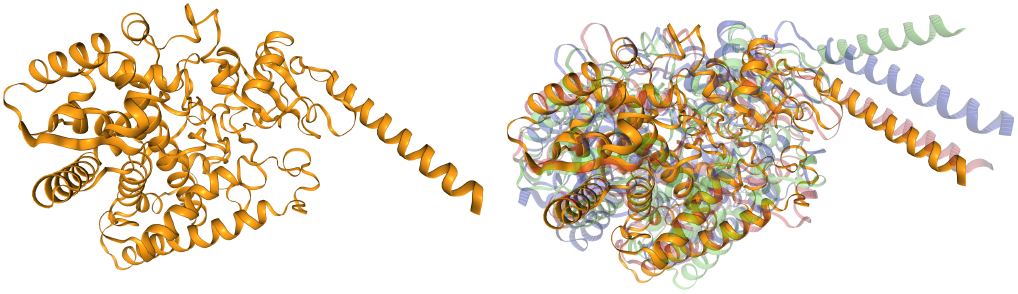
Cytochrome P450 family 76 subfamily C polypeptide 7 from *Zea mays* (UniProt ID: A0A1D6JW22) in yellow. Examples of AlphaFind search results: Uncharacterized protein from *Oreochromis niloticus* (UniProt ID: A0A669CAC6) in blue, Cytochrome P450 2C19 from *Equus caballus* (UniProt ID: A0A3Q2H4N0) in green, Cytochrome P450 2H1 from *Haliaeetus albicilla* (UniProt ID: A0A091PH09) in red.

The PIN proteins are transmembrane proteins which regulate plant growth by influencing auxin transport from the cytosol to the extracellular space. They only occur in plants and feature a configuration of 10 main helices that collectively form a pore. Eight types of PIN proteins are known (PIN1-PIN8), and recently, the structures of three PIN proteins were uncovered and published in Nature. The structure of the PIN5 protein differs from other PINs (17) and has not yet been experimentally determined. This use case shows (Figure 4) that the PIN5 protein structure is strongly conserved among many different plant species.

**Figure 4.**
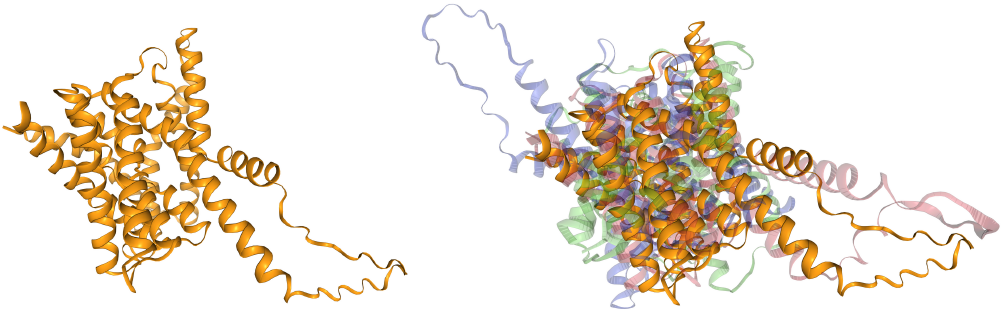
Auxin efflux carrier component 5 from Arabidopsis thaliana (UniProt ID: Q9FFD0) in yellow. Examples of AlphaFind search results: *Auxin efflux* carrier component from *Citrus unshiu* (UniProt ID: A0A2H5NWV3) in blue, uncharacterized protein from *Eucalyptus grandis* (UniProt ID: A0A059BG64) in green, Auxin efflux carrier component from *Vitis vinifera* (UniProt ID: A0A438CTF0) in red.

## CONCLUSION

In this article, we presented AlphaFind, a novel web application for fast structure-based search of similar proteins in AlphaFold DB and PDB. AlphaFind utilises the Learned Metric Index approach and a 3D feature extraction method designed for protein structures. The web application presents the search results as a table, sortable according various criteria, i.e., TM-score, RMSD, and the number of aligned residues. The users can also download the results as a CSV file or visualize superpositions of input and output proteins via NGL Viewer. AlphaFind is easy to use and is platform-independent. Documentation describing its usage is referenced on the web page https://alphafind.fi.muni.cz.

## DATA AVAILABILITY

AlphaFind application is available at no cost and no registration at https://alphafind.fi.muni.cz. The user manual is referenced in the application and is also directly available at https://github.com/Coda-Research-Group/AlphaFind/wiki/Manual. The application’s source code is accessible under the MIT license at https://github.com/Coda-Research-Group/AlphaFind and is also available in the Supplementary Data. The LMI is published and its performance reproducibly examined in (18). The embeddings and weights for the LMI model are available in the Czech National Repository’s pilot at https://doi.org/10.48700/datst.d35zf-1ja47.

## SUPPLEMENTARY DATA

Supplementary data are available at NAR Online.

## ACKNOWLEDGEMENTS

The authors would like to thank Lucie Novotná for her help with the protein embeddings, Lukáš Hejtmánek for his work on the container infrastructure, and Vladimír Míč and Tomáš Raček for consultation during the early stages of our involvement with the protein structure data.

